# PALADIN:Protein Alignment for Functional Profiling Whole Metagenome Shotgun Data

**DOI:** 10.1101/047712

**Authors:** Anthony Westbrook, Jordan Ramsdell, Taruna Aggarwal, Louisa Normington, R. Daniel Bergeron, W. Kelley Thomas, Matthew MacManes

**Affiliations:** University of New Hampshire, Department of Computer Science; University of New Hampshire, Hubbard Center for Genome Studies; University of New Hampshire, Department of Molecular Cellular and Biomedical Sciences

## Abstract

Whole metagenome shotgun sequencing is a powerful approach for assaying the functional potential of microbial communities. Presently, we lack tools that efficiently and accurately align DNA reads against protein references, the technique necessary for constructing a functional profile. Here, we present PALADIN—a novel modification of Burrows-Wheeler Aligner that provides more accurate alignment and orders-of-magnitude improved efficiency by directly mapping in protein space.

---

As high-throughput sequencing technologies improve, the analysis of microbial community composition and function has rapidly advanced. Historically, this has mostly focused on taxonomic surveys using a small number of phylogenetically informative genes such as the small subunit of ribosomal RNA^1,2^. The ability to taxonomically profile communities provided new insights into the role of microbiomes in human health^3,4^, soil ecology^5,6^ and environmental remediation^7^. Nevertheless, the gene survey approach provides limited functional knowledge because microorganisms with similar or even identical rRNA sequences often differ significantly with respect to genomic content, and therefore may have vastly different functional roles in their environment^8^.

Functional profiling of microbial communities based on Whole Metagenome Shotgun (WMS) sequencing data attempts to catalog the genes present in a community. An inventory of the protein coding functions of a microbial community can be created by either matching the individual reads to annotated reference databases or by assembling the reads and annotating the resulting chromosomal segments^9^. Conventional methods such as BLAST are robust but computationally intensive and techniques for rapidly mapping DNA reads to annotated reference genes fail when the references within the curated databases diverge moderately from DNA sequences of homologous genes in the metagenome sample. To mitigate these challenges, researchers often turn to metagenome assembly and subsequent annotation which has profound shortcomings, such as chimeric assembly of closely related sequences, strong bias toward abundant organisms, and substantial human and computer resource requirements^2,9^. Therefore, current approaches are not sufficient to satisfy the requirements of researchers attempting to understand functional metagenomics.

To improve the sensitivity with which functional profiling of metagenomics samples is performed, we present PALADIN—an algorithm adapted from the popular BWA^11^ mapping tool (source code is available at https://github.com/twestbrookunh/paladin, see supplementary note 4 and figure 5 for implementation details). In brief, PALADIN identifies and translates six possible open reading frames within each read, and maps these translated DNA sequence reads to a protein reference allowing for rapid identification of functional metagenomic profiles. By mapping in protein space, this method takes advantage of the general conservation of amino acid sequences compared to the underlying DNA sequences. Here we demonstrate the application of this modified alignment algorithm for rapid and sensitive functional metagenomic profiling using large scale WMS datasets. PALADIN reports mappings in standard SAM format, and can generate a tab-delimited file from which additional information can be obtained, including protein abundance and gene ontology.

To evaluate the performance of this novel protein space read mapper, we first generated typical 250 basepair long paired-end reads using the standard Illumina error model for six well-annotated bacterial genomes *(Pseudomonas fluorescens, Escherichia coli, Acidovorax avenae, Micrococcus luteus, Halobacillus halophilus* and *Staphylococcus epidermidis)* using the read simulation package ART^12^. The reads for the six genomes were pooled to create a mock-metagenomic read dataset (see supplementary note 1). To establish a positive control, we used PALADIN to map the combined reads to the protein sequences of the six original genomes and to a filtered Swiss-Prot database (see supplementary note 2), which contains well-curated protein sequences excluding the bacteria from which the reads were derived. To compare PALADIN’s relative mapping efficiency and accuracy to existing tools that align reads only in DNA space, we mapped the nucleotide version of the reads to two reference types using BWA and Novoalign (http://www.novocraft.com)—a DNA based read mapper with the capacity to map reads to a degenerate nucleotide reference (see supplementary note 3). As expected, when mapping reads to the genomes they were derived from, PALADIN and BWA map 98.29% and 96.02% of these reads respectively; By Contrast, Novoalign implementing the degenerate read mapping methodology mapped 36.39% of the reads. The poor performance of Novoalign appears to stem from the fact that while it accepts degenerate bases in the references it does not score them as positive matches during alignment.

By leveraging our prior knowledge about the genes corresponding to the simulated reads, we evaluated the functional accuracy of mapping using three metrics—percentage of reads mapped, Jaccard similarity coefficient^13^, and the number of unique proteins found in Swiss-Prot. To calculate the similarity coefficient, reads and their corresponding aligned targets were assigned functionality using the standardized Gene Ontology (GO) language (see supplementary note 5). Each GO term represents a vertex within a graph formation where conditional edges join terms tracing back to one of three parent vertices: biological processes, molecular function, and cellular component. For each read and its matching Swiss-Prot entry, graphs were constructed by the GO term assignments directed back to their respective parent vertices. The ratio of the intersection and the union of both graphs was used to determine the Jaccard similarity coefficient as an alignment accuracy metric. The number of unique proteins was determined based on each distinct Swiss-Prot ID the reads mapped to.

When we tested the ability of PALADIN to map mock-metagenomic reads to the Swiss-Prot database, it mapped 30.65% of compared to 19.79% (BWA) and 0.56% (Novoalign), while all three systems mapped with extremely high accuracy (correct functional assignment) measured by the similarity index (Table 1). These results suggest that mapping in protein space as implemented in PALADIN is even more accurate than existing solutions while using a larger proportion of the available shotgun metagenomics dataset.

**Table 1.**
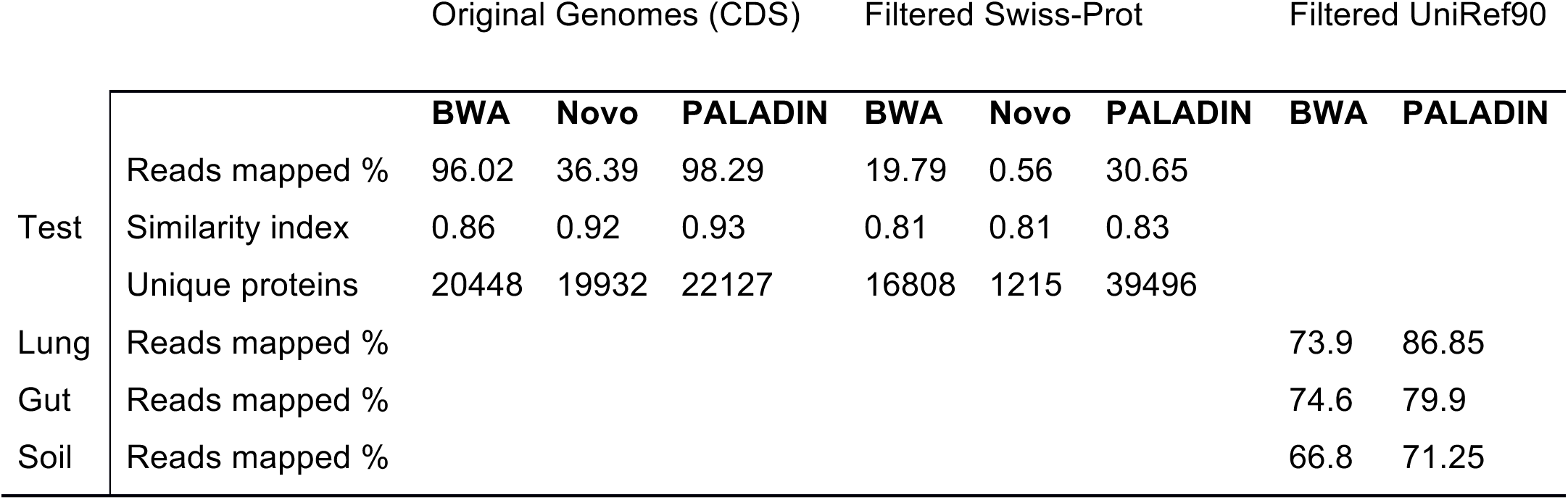
Mapping efficiency of PALADIN, BWA, and Novoalign against multiple read sets. Positive control was established by aligning the simulated reads against the original six test genomes. This simulated read set was then aligned against the filtered Swiss-Prot reference, resulting in PALADIN performing best in percentage of reads mapped, similarity index, and number of aligned proteins. Additionally, three empirically derived read sets (lung (BioProject:PRJNA71831), gut (BioSample:SAMN00037421), and soil (MG-RAST:4520320.3)) were aligned against the filtered UniRef90 reference, each resulting in PALADIN scoring better than both BWA and Novoalign.

While mapping mock reads to the well annotated SwissProt database allows us to assay accuracy via the use of GO-term similarity, it is a relatively small dataset with limited representation of both functional and taxonomic breadth. A more ideal reference would be the UniRef90 database which contains taxonomically diverse sequences clustered at 90% sequence identity. In a process identical to above, we mapped translated DNA reads from three published WMS dataset from different environments to the Uniref90 proteins using PALADIN, and untranslated reads to the corresponding nucleotide sequences from Uniref90 proteins using BWA. With mapping accuracy established in the first set of experiments, the enhanced mapping rate using PALADIN versus BWA (Table 1) translates to increased resolution of functional profiling.

To contrast the differences in computational efficiency between PALADIN and conventional protein alignment tools, we mapped the reads of a dataset consisting of nearly 240,000,000 reads against the Uniref90 database with PALADIN using 28 cores on a high-end workstation. The computation finished after 31 hours for an efficiency of approximately 128,000 reads/minute. We then extracted 8,000 of these sequences from the dataset and performed an alignment with BLAST^10^ using the same hardware environment and resource availability. BLAST completed execution in 8.5 hours with an efficiency of about 16 reads/minute. Given the linear time complexity associated with BLAST, we estimate that an execution run against the full dataset would take about 29 years, approximately 8,000 times longer than PALADIN.

In summary, we present PALADIN, a tool for accurate functional characterization of metagenomic samples that is orders of magnitude faster than existing approaches. This significant improvement in efficiency affords researchers unprecedented opportunity to gain detailed and novel insight into microbial communities. Additionally, by constructing this approach upon a widely used alignment algorithm, reliability and usability are inherently increased, which promotes faster adoption and easier incorporation into existing pipelines. Finally, the reduction in required computational resources creates a more cost-effective solution, thereby increasing viability of analysis capabilities in environments where economic pressures are present. Given these aspects, PALADIN may potentially aid in any number of evolving fields that depend on functional characterization, including personalized medicine, biodefense, environmental remediation, transcriptomics, and the study of emerging pathogens.

